# Distinct gaits of self-propelled quadriflagellate microswimmers

**DOI:** 10.1101/2022.05.10.491287

**Authors:** Dario Cortese, Kirsty Y. Wan

## Abstract

Legged animals often coordinate multiple appendages for both underwater and terrestrial locomotion. Quadrupeds in particular, change their limb movements dynamically to achieve a number of gaits, such as the gallop, trot, and pronk. Surprisingly, micron-sized unicellular algae are also capable of coordinating four flagella to produce microscale versions of these gaits for swimming. Here we present a fully-3D model of a quadriflagellate microswimmer comprising five beads and systematically investigate the effect of gait on swimming dynamics, propulsion speed, efficiency, and induced flow patterns. We find that by changing gait alone, distinct motility patterns emerge from the same basic microswimmer design. Our findings suggest that different species of morphologically-similar microorganisms (e.g. with identical number and placement of appendages) evolved distinct flagellar coordination patterns as a consequence of different ecological drivers. By comparing the flagella-induced flows in terms of volumetric clearance rate, we further explore the implications of distinct gaits for single-cell dispersal, feeding, and predator-avoidance.

## I. INTRODUCTION

Biological microswimmers display a variety of species-specific locomotor behaviours. In motile unicellular algae, diverse ciliary or flagellar beat patterns result in different microscale swimming gaits, some of which are comparable to those of animals with analogous limb positioning. A single species can perform multiple swimming gaits depending on environmental conditions, or even switch dynamically between them. For example, the model organism *Chlamydomonas* swims by beating its two flagella in a quasi-synchronous breaststroke pattern; however, losses of synchrony can stochastically trigger a transient second gait (‘phase slip’) in which flagella beat out-of-phase [1, 2]. Quadriflagellates, algae bearing four flagella, are another commonly-occurring configuration of microswimmer, abundant in many freshwater and marine habitats [3–5]. Their four appendages can have equal lengths and beat patterns, or they may have different lengths, beat patterns, and functions, depending on the stage of basal-body maturation [6]. It has been shown recently that quadriflagellate gaits can be classified by comparing them to the movement of quadrupeds [7]. Gaits such as the trot, pronk, gallop have all been observed, in which the relative phase relationships between the beating flagella can be confirmed by high-speed imaging [8].

At the microscale, what is the rationale behind selecting a specific gait in a given situation, and what are the advantages and drawbacks of individual gaits? Moreover, why do different species preferentially select for one gait over another? The answer deviates significantly from our intuitions associated with macroscopic vertebrate locomotion [9]. For instance, *Pyramimonas parkeae* swims using a ‘trot’ gait, achieving an average speed of up to 400 *µ*m*/s*, several times faster than its sister species *Pyramimonas tetrarhynchus*, which largely swims using the ‘pronk’ gait [7, 10]. In terms of cell size and shape, both species comprise obovate or oblong cells of approximately 20 *µ*m in length, 10 *µ*m in width, and four equal-length, front-mounted flagella (10 *−*15 *µ*m), arranged in a cruciate pattern (Fig. 1A). Despite their near-identical body and flagella morphology, different quadriflagellate swimmers nonetheless exhibit distinct motility characteristics [10]. Therefore, we ask if and how gait alone can influence swimming dynamics.

**FIG. 1.**
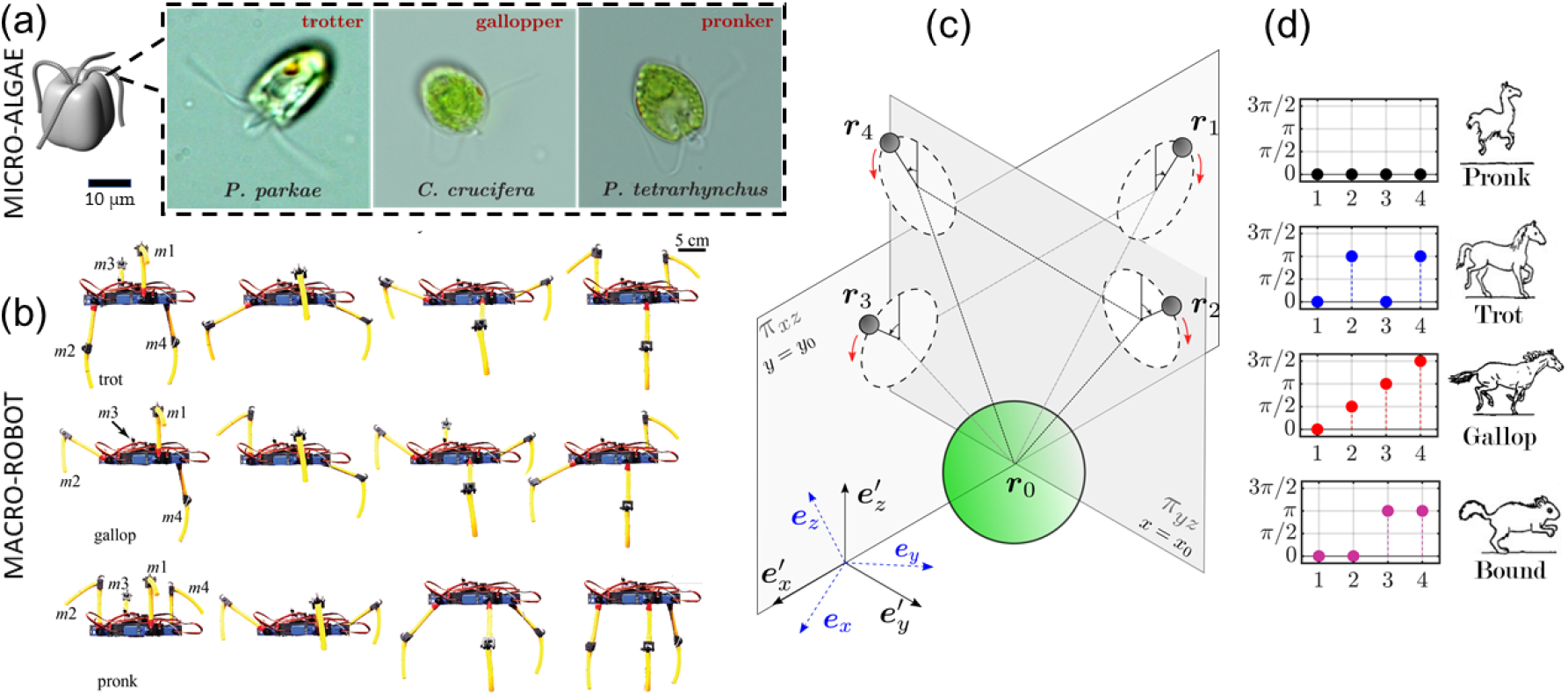
(a) A stereotypical quadriflagellate cell has four front-mounted flagella (cilia) emanating from an anterior grove, where they assume a cruciate configuration. Different species of these algae assume distinct swimming gaits including the “pronk”, “gallop”, and “trot”. (b) A macroscopic robot modelled on these algae can achieve different rates of self-propulsion at low-Reynolds number, depending on gait (image modified from [10]). (c) Here, we model a quadriflagellate microswimmer using a system of 5-beads, and systematically evaluate the swimming performance of each gait. The flagella beads rotate on circular orbits embedded in independent planes that may be tilted (the zero-tilt case is shown for simplicity). (d) Representations of these gaits in legged-animals.

Any microswimmer’s ability to swim depends strongly on its intrinsic shape and sequence of shape deformations. Many simple microswimmer designs have been studied extensively using theoreticalor computational approaches, such as Purcell’s three-link swimmer [11], the linear three-bead Najafi-Golestanian swimmer [12], the two-sphere pushme-pullyou swimmer [13], a *Chlamydomonas-like* planar triangular swimmer [14], and many others. Another swimmer composed of a I-shaped frame and four rotating disks (termed Quadroar), was shown to be capable of 3D maneouvreability [15]. Unlike in experiments, simulation parameters can be readily varied to evaluate how shape and gait influence propulsion efficacy, or to identify the existence of optimal gaits [16]. To date, only some of these systems have been realised experimentally, for example using colloids or beads that are driven by optical tweezers or external magnetic fields [17–19]. In-depth studies of such analytically tractable microswimmers will not only help us understand the biological propulsion mechanisms of extant microorganisms, but also inform and improve the design and control of artificial or robotic swimmers that are capable of effective navigation at low-Reynolds numbers.

In this article, we systematically explore the dependence of microswimming on gait by developing a novel *in silico* model of a quadriflagellate alga (Fig. 1). We compare a range of quadriflagellate gaits by solving the appropriate zero-Reynolds number hydrodynamics equations, and compute the resulting trajectories and both the near and far-field flow patterns.

We note that when path-sampling the trajectory history of live cells, stochastic transitions between different modes or gaits can occur due to biological noise and environmental perturbations, and this can significantly impact the overall swimming trajectory [20–22]. For simplicity, we shall neglect gait-switching dynamics here and focus only on the consequences of a sustained, deterministic gait. We will also discuss our results in light of available biological data, as well as measurements from a recent macroscopic (𝒪 (10)cm) robophysical model ([10]) of quadriflagellate algae.

The paper is organised as follows. We begin by introducing the quadriflagellate model, and associated equations of motion (Section II). We then compare the three-dimensional flow fields produced by such a swimmer, for a variety of prescribed flagellar actuation gaits (Section III). In Section IV, we consider the impact of these flows on the free swimming trajectories, and extend the model to explore the role of asymmetries in the either the flagellar geometry or the beat frequency (Section V). Finally, in Section VI, we explore possible functional relationships between gait and induced flows for processes unrelated to swimming, such as enhanced feeding, fluid mixing, or cloaking from predators.

## II. MODEL FORMULATION

Our model quadriflagellate consists of five beads immersed in an incompressible Newtonian fluid, interacting hydrodynamically at zero-Reynolds number. The cell body has radius *a*_0_ and is located at ***r***_0_. The four flagella are modelled by smaller beads of radius *a* ≪ *a*_0_, located at the centers of drag of each flagellum ***r***_*i*_, *i* = 1 *−* 4. These are constrained to move along circular orbits of radius *R*. The 4 orbits are fixed on a rigid triangular scaffold at rest with the body frame of reference (Fig. 1c), with dimensions *𝓁* and *h*, and *a* ≪ *a*_0_ ≪ *h, 𝓁*. Similar orbiting-bead models have been used to investigate hydrodynamic synchronisation of beating cilia in planar configurations [23–26]. More recently, a three-dimensional, 3-bead version of this model helped reveal the origins of the superhelical swimming motion typical of the biflagellate alga *Chlamydomonas reinhardtii* [27]. The underlying motivation for such models comes from experimental evidence that flow-fields induced by freely-swimming *Chlamydomonas* cells are well-described by just three Stokeslets [28].

Similar to the biflagellate case, here we also allow for out-of-plane swimming by tilting the single-flagella orbital planes - here, flagellar beads produce the axial rotation needed for helical swimming (Fig. 11, inset). Each flagellar bead is associated with an independent tilt angle *β*_*i*_, *i* = 1 *−* 4. All four flagella have the same tilt, unless otherwise stated (e.g. Fig. 5).

The beads are assumed to move along circular orbits centred at

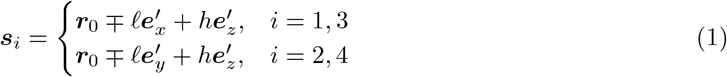

When only two flagella are present (say flagella 1 and 3, and *β*_1_ = *β*_3_ = 0), we recover the familiar in-plane breaststroke configuration.

The flagellar beads are located at ***r***_*i*_ = ***s***_*i*_ + *R****n***_*i*_ with

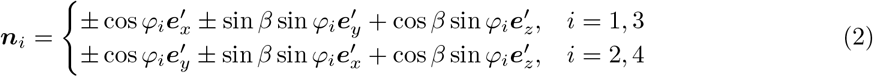

The body axes 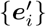 transform to the lab frame {***e***_*i*_} via Euler angles ***θ*** = (*θ*_1_, *θ*_2_, *θ*_3_) (respectively, yaw, pitch and roll).

In many types of real and artificial cilia, the stroke speed is not perfectly uniform throughout the beat cycle [29, 30]. In order to implement a variable beat cycle along a perfectly circular orbit we allow the frequency to vary slightly during the cycle to mimic the asymmetric stroke pattern observed in most protist cilia. The inclusion of such a nonlinearity also ensures that the average stress pattern exerted by the cells on the fluid as measured in experiments is reproduced [28, 31]. This can be achieved by asymmetric force profiles [27, 32], or asymmetric (such as elliptical) trajectories [33].

Here, we set the beads to rotate with nonlinear phase dynamics described by

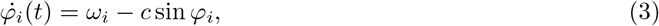

and beat frequency 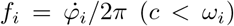. The four flagella beat with the same frequency, unless otherwise indicated. We note that in contrast to a previous formulation [27], the model quadriflagellate microswimmer is prescribed by the imposed gait kinematics, rather than by the tangential component of the flagellar forces.

Once the flagellar beating pattern *φ*_*i*_(*t*) is set, the swimmer kinematics are fully described by the vector (*x*_0_, *y*_0_, *z*_0_, *θ*_1_, *θ*_2_, *θ*_3_, *φ*_1_, *φ*_2_, *φ*_3_, *φ*_4_) and the parameters *h, 𝓁, β, R, a*_0_, *a*. The swimmer has translational velocity 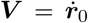 and angular velocity **Ω**. The governing equations of the model can be derived from the force balance on each flagellar bead. Each flagellum is driven by a tangential force 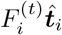 and it is balanced by the tangential component of the local hydrodynamic drag (given by Stokes’ law), 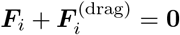:

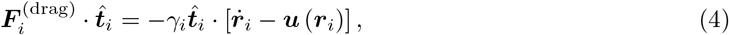

where the dot denotes a time derivative, *γ*_*i*_ = 6*πηa*_*i*_ and *η* is the dynamic viscosity of the fluid. Because each bead is constrained to move on a circular trajectory (in the frame of the swimmer), the tangential force balance becomes

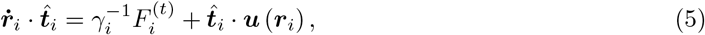

where the velocity flow field ***u*** satisfies Stokes equation and the incompressibility condition:

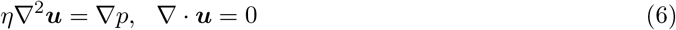

If we consider each sphere in the far field of the others, the flow field at any point can be expressed, at first order, using the Oseen approximation for hydrodynamic interactions between beads:

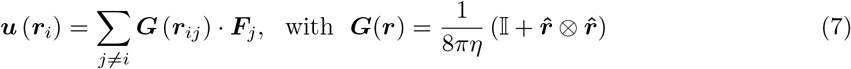

where ***r***_*ij*_ = ***r***_*i*_ *−****r***_*j*_. Similar relationships apply to the rotational motion of each bead. The dynamics of the model are therefore governed by the following set of equations:

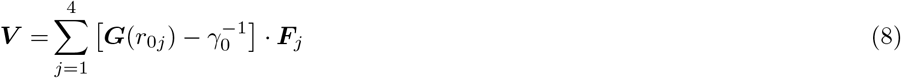

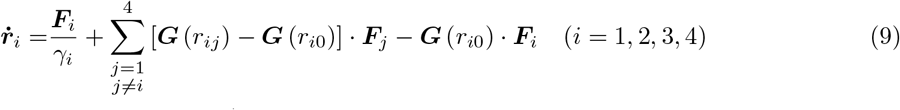

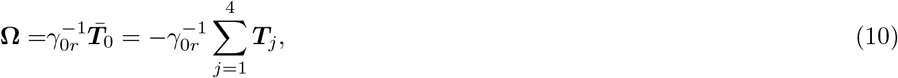

where 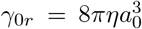. ***T***_*i*_ are the torques in the laboratory frame of reference, and 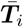 denotes the intrinsic torques due to the rotation of the beads around an internal axis; *a*_0_ ≫ *a* implies 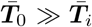. The velocity of each bead is linked to the phase dynamics via 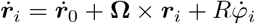, thus coupling equation (10) to (8)-(9).

We impose force-and torque-free conditions:

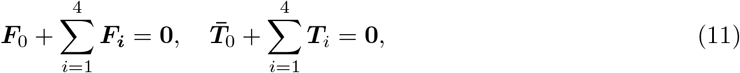

where 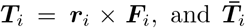 is the intrinsic torque due to the *i*th sphere’s rotation around an internal axis. Equations (9)-(10) reduce to a set of 14 equations for unknowns (***V***, **Ω, *F***_*i*_). The system is solved numerically by an iterative process. First, morphological parameters are chosen to match the geometrical properties of a typical quadriflagellate. All lengths are non-dimensionalised by *𝓁* (= 10 *µm*, typical cell size), forces by the average tangential flagellar force *F*_0_ (= 30 pN, typical force produced by a flagellum), and *η* by 10^*−*3^ pN*µm*^*−*2^ (viscosity of water) [27]. Second, the boundary and initial conditions are chosen, including the parameters determining the gait, *c* and *ω*_*i*_, and the initial velocities ***V*** (*t* = 0), Ω(*t* = 0), forces and positions ***r***_*i*_, ***θ***. The equations are then solved for the velocities and forces (***V***, **Ω, *F***_*i*_); these are in turn used to propagate the positions to the following time step. A fixed time step was used. The method of quaternions was used to resolve singularities at *θ*_2_ = *±π/*2 [34] (see also [27]).

## III. FLUID FLOWS

We used the above model to explore 4 distinct gaits of quadriflagellate self-propulsion, namely pronk, trot, bound and gallop. We assume a fixed geometry: *a*_0_ = 0.7, *a*_1*−*4_ = 0.05, *h* = 1.3, *𝓁* = 1, *R* = 0.6 and *β*_1*−*4_ = 0.3, and only vary the gait - via the flagellar phases. Table 1 summarises the phase relationships between the four flagella in each of the different gaits (see also Supplementary videos 1). We considered two variants of the gallop gait, clockwise, or respectively counter-clockwise.

**Table I.** List of the corresponding phase relationships for each of the tested quadriflagellate gaits, relative to the phase of flagellum 1 (*φ*_1_). The final column corresponds to the two-flagella case, which resembles the in-phase breaststroke of *Chlamydomonas reinhardtii* (CR).

| $\varphi_1$ | Pronk | Trot | Gallop (CW) | Gallop (CCW) | Bound | CR |
| --- | --- | --- | --- | --- | --- | --- |
| 0 | (0, 0, 0, 0) | (0, $\pi$, 0, $\pi$) | (0, $\frac{\pi}{2}$, $\pi$, $\frac{3}{2}\pi$) | ($\frac{3}{2}\pi$, $\pi$, $\frac{\pi}{2}$, 0) | (0, 0, $\pi$, $\pi$) | (0, -, 0, -) |
| $\frac{\pi}{2}$ | ($\frac{\pi}{2}$, $\frac{\pi}{2}$, $\frac{\pi}{2}$, $\frac{\pi}{2}$) | ($\frac{\pi}{2}$, $\frac{3}{2}\pi$, $\frac{\pi}{2}$, $\frac{3}{2}\pi$) | ($\frac{\pi}{2}$, $\pi$, $\frac{3}{2}\pi$, $2\pi$) | ($2\pi$, $\frac{3}{2}\pi$, $\pi$, $\frac{\pi}{2}$) | ($\frac{\pi}{2}$, $\frac{\pi}{2}$, $\frac{3}{2}\pi$, $\frac{3}{2}\pi$) | ($\frac{\pi}{2}$, -, $\frac{\pi}{2}$, -) |
| $\pi$ | ($\pi$, $\pi$, $\pi$, $\pi$) | ($\pi$, $2\pi$, $\pi$, $2\pi$) | ($\pi$, $\frac{3}{2}\pi$, $2\pi$, $\frac{\pi}{2}$) | ($\frac{\pi}{2}$, $2\pi$, $\frac{3}{2}\pi$, $\pi$) | ($\pi$, $\pi$, $2\pi$, $2\pi$) | ($\pi$, -, $\pi$, -) |
| $\frac{3}{2}\pi$ | ($\frac{3}{2}\pi$, $\frac{3}{2}\pi$, $\frac{3}{2}\pi$, $\frac{3}{2}\pi$) | ($\frac{3}{2}\pi$, $\frac{\pi}{2}$, $\frac{3}{2}\pi$, $\frac{\pi}{2}$) | ($\frac{3}{2}\pi$, $2\pi$, $\frac{\pi}{2}$, $\pi$) | ($\pi$, $\frac{\pi}{2}$, $2\pi$, $\frac{3}{2}\pi$) | ($\frac{3}{2}\pi$, $\frac{3}{2}\pi$, $\frac{\pi}{2}$, $\frac{\pi}{2}$) | ($\frac{3}{2}\pi$, -, $\frac{3}{2}\pi$, -) |
| $2\pi$ | ($2\pi$, $2\pi$, $2\pi$, $2\pi$) | ($2\pi$, $\pi$, $2\pi$, $\pi$) | ($2\pi$, $\frac{\pi}{2}$, $\pi$, $\frac{3}{2}\pi$) | ($\frac{3}{2}\pi$, $\pi$, $\frac{\pi}{2}$, $2\pi$) | ($2\pi$, $2\pi$, $\pi$, $\pi$) | ($2\pi$, -, $2\pi$, -) |

With reference to biological microswimmers, the pronk, trot, and gallop gaits are adopted routinely by different quadriflagellate species [7, 35]. Meanwhile the bound gait has only been observed occasionally, including in quadriflagellate zygotes of *C. reinhardtii* immediately after mating and cell-cell fusion [7].

As the model quadriflagellate swims, it displaces the surrounding fluid and exerts stresses that induce specific flow patterns. The instantaneous fluid flow velocity field ***u***(***r***, *t*) can be calculated from (7) as

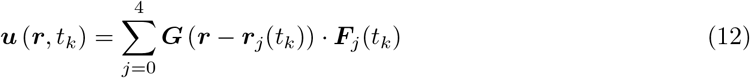

by superimposing the 5 stokeslet fields located at the positions ***r***_*j*_(*t*_*k*_), obtained by solving (8)-(10) at each time step *t*_*k*_ = *k* Δ*t*. Beat-cycle averages can be derived by calculating

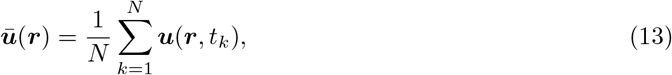

where *N* = (*f* Δ*t*)^*−*1^ is the number of time steps in a beat cycle.

The resulting flow field is three-dimensional and characterised by gait-dependent flow features. Fig. 2 shows the evolution of the velocity vector field in the fluid at five different stages of the beat cycle, for four distinct gaits. Since the tilt angle *β* ≠ 0 (indicating a non-planar tilt angle for all four flagella), there is an intrinsic chirality for all gaits. Here we have chosen *β*_1*−*4_ = 0.3 based on experimental measurements obtained from *C. reinhardtii* [27] (similar measurements have not yet been obtained for quadriflagellate species); this parameter can be easily changed. For gaits exhibiting a high-degree of rotational symmetry, such as the pronk or trot, the flow field is highly symmetrical with respect to the body axis, whereas a net asymmetry is observed in the flow patterns ensuing from bound and gallop gaits. In the CW-gallop, a small vortex can also be seen to propagate in the clockwise sense. The flow fields also display novel flow features not seen around standard pushers, pullers or squirmers (see Supplementary videos 2-5).

**FIG. 2.**
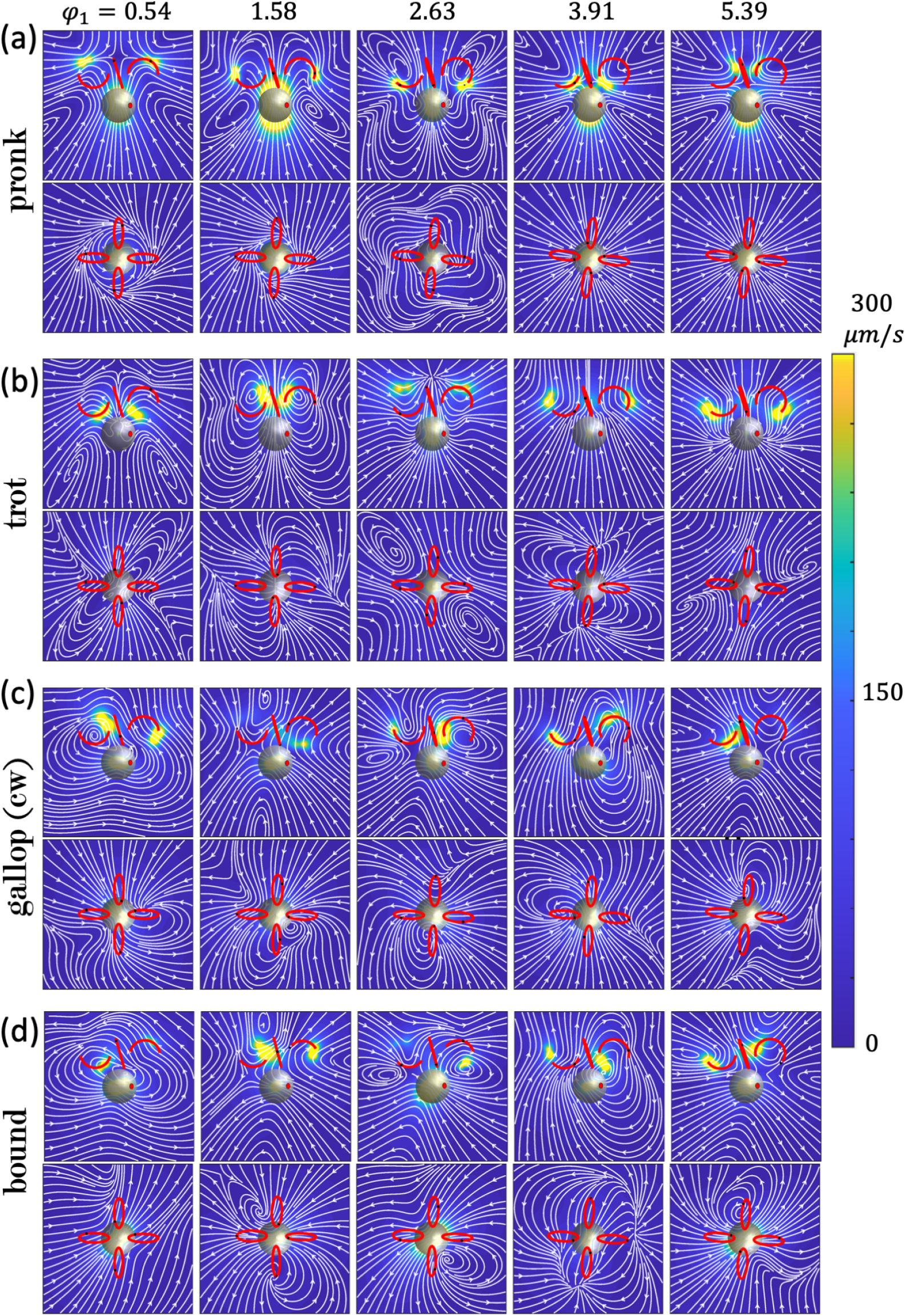
Time dependent flow fields around a quadriflagellate microswimmer performing four distinct gaits pronk, (b) trot, (c) gallop (CW), and (d) bound, in each case as observed in horizontal slices in the xz (side-view) and xy (top-view) profiles (B).

The periodic alternation between power and recovery strokes leads to characteristic oscillatory flow patterns in the far-field (Fig. 3a). During the recovery stroke phase of each beat cycle, the flow field is that typical of a pusher swimmer - the fluid is pushed out from the sides, and drawn inwards from above and below, while the pattern typical of a puller swimmer is observed during the power stroke. A similarly time-varying, oscillatory flow pattern has also been observed previously in simulations of a planar 2-flagella swimmer [36]. Moreover, the time-averaged flow fields display the characteristic profile of a puller swimmer, similar to that of the biflagellate *Chlamydomonas*. For brevity, only the trot flow field is shown in Fig. 3b,c. For all gaits, the cycle-averaged flow field displays a stagnation point in front of the flagellar orbits, again similar to the biflagellate case.

**FIG. 3.**
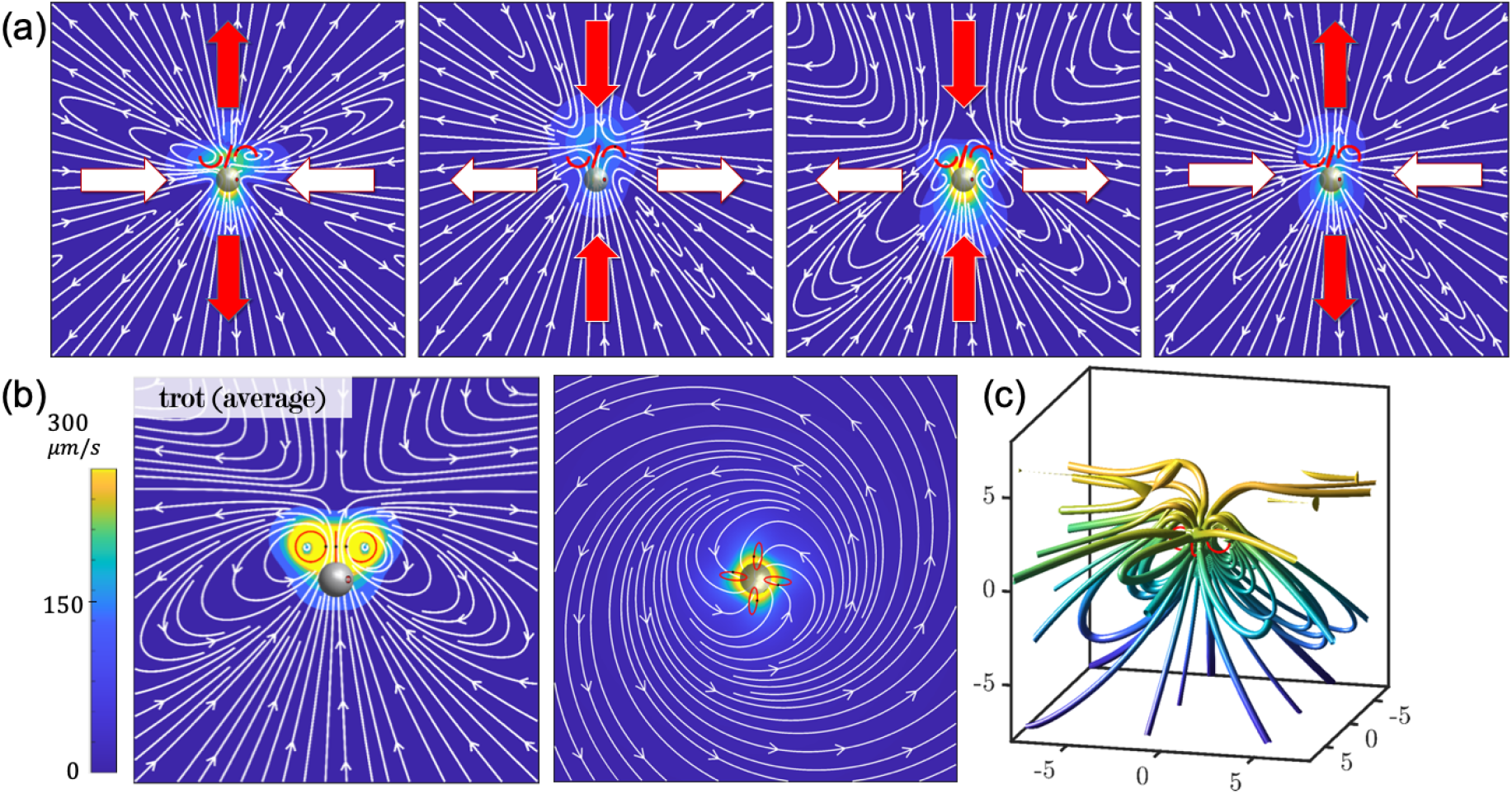
(a) Far-field flow patterns for the trot gait alternates between pusher and puller flow fields over the course of a complete beat cycle. For the trot gait, the time-averaged flow fields are viewed from the side and top (b), together with a 3D rendering (c). (Note: (c) is plotted in computational units.)

## IV. SWIMMING SPEED AND ROTATION

Flagellated algae, like several other swimming microorganisms, assume superhelical swimming trajectories. Superhelices are three-dimensional curves characterised by small-scale, fast swirls modulated by a larger-scale, slow helix. Such trajectories have been identified as the general solutions to the swimming problem at low Reynolds numbers [37]. Superhelices have been observed in sperm swimming [38] and in biaxial self-propelled particles under external torques [39].

Superhelical motion results from two concurrent rotations around two perpendicular axes. In our previous study on biflagellate swimmers, we showed that the superhelical trajectories typical of *C. rehinardtii* can be reproduced by a three-dimensional 3-bead model with non-planar beat dynamics and a small (*>* 1%) asymmetry in the forces driving the individual flagella. The bilateral asymmetry occurs simultaneously with a persistent axial rotation. While swimming forward with velocity ***V***, the alga rotates around its body axis with angular velocity **Ω**. This rotation results from the non-planarity of the flagellar beat, which generates a net torque. Thus, fast swirls arise as a consequence of the alternation of non-planar power and recovery strokes which make up the full cycle of a typical flagellar beat. On the other hand, the large-scale helical character of the trajectories is the result of the asymmetric motion of the two flagella [27].

The free-swimming trajectories induced by the 5-bead swimmer model according to (9)-(10) are also superhelical (see Fig. 4). We introduce an orbital tilt *β*_*i*_ = 0.3 to allow for beat non-planarity. Different gaits are implemented according to Table 1, by imposing an initial condition on the phases 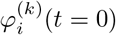, with *k* = pronk, trot, gallop, bound, half-trot, and *i* = 1 *−* 4.

**FIG. 4.**
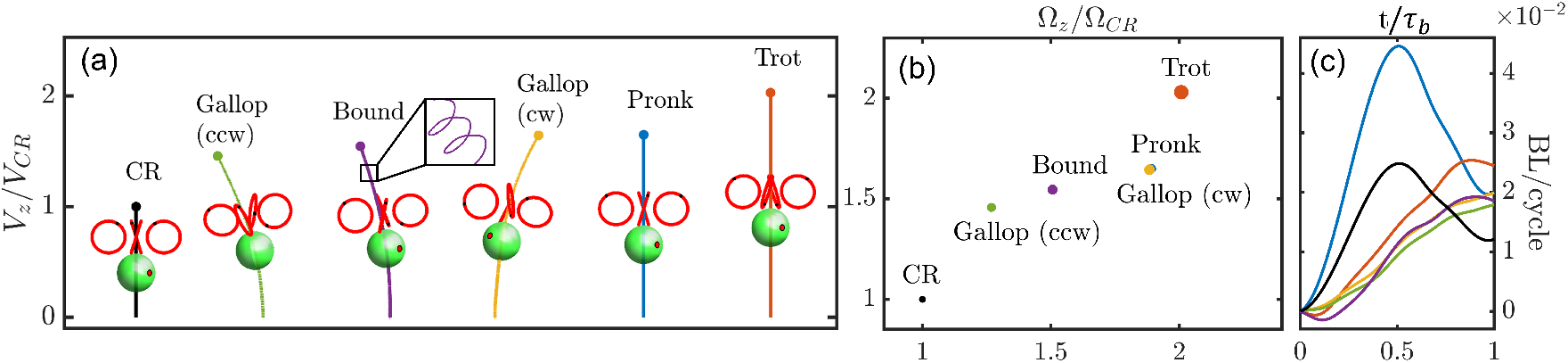
(a) Similar to a biflagellate (CR) executing an in-phase breaststroke, a model quadriflagellate microswimmer traces out assorted superhelical trajectories that depend on gait. Inset: fast in-beat swirls occurring on the timescale of ciliary beating. (b) Scatter plot of the translational and rotational speeds of the different gaits. Marker size is scaled by the respective Lighthill efficiency *ε* of each gait (see text). (c) The within-beat cycle displacement (relative to the starting position) is plotted against the scaled time *t/τ*_*b*_, where *τ*_*b*_ is the beat period of the corresponding gait.

We found that the average net linear and rotational velocities achieved by the model swimmer in different gaits vary considerably. For a fixed body geometry and *β*_*i*_ = 0.3 for the single-flagellum tilt angle, Fig 4 shows the angular velocity Ω and the forward speed *V*_*z*_ obtained in simulations for each of the gaits we tested. Average net speed was calculated by averaging over 20 beat cycles, with 100 time steps per beat cycle. We observed that cells undergoing more symmetric gaits such as the biflagellate breaststroke (CR), and the quadriflagellate pronk and trot, proceed smoothly forward along perfectly straight trajectories Meanwhile more bilaterally-asymmetric gaits such as the bound and gallop produce more curved trajectories, where the overall axis-of-progression of the superhelices are tilted. In the case of the gallop, the sense of gait rotation also dictates the overall helix tilt direction.

We further evaluated the rotation-translation coupling for the different gaits, and compared their swimming performance against our two-flagella reference - modelled on *Chlamydomonas* (CR). The ‘trot’ gait produced the fastest linear progression as well as the fastest angular progression. All other quadriflagellate gaits result in largely similar linear speeds, but segregated in terms of angular rotation speed (Figure 4b), all falling between 1 *−* 2 times that for the biflagellate CR.

These gait-dependent linear swimming speeds are consistent with the available data, in which the trot gait proceeds at 408 *µ*m/s, much faster than the pronk at 126 *µ*m/s, and gallop at 127 *µ*m/s [10], and several times faster than that of CR at *∼*90 *µ*m/s [40, 41]. Due to technical limitations in visualising the free-swimming trajectories in 3D, experimental estimates of the axial rotation speeds of the quadriflagellate species are not yet available. Our results are in broad agreement with measurements using an upscaled robophysical model in which the four flagella are actuated according to different gaits. In this case the mechanical flagella are configured relative to the cell body axis without tilt (*β* = 0). The robot experiments (Fig. 1B) revealed a similarly strong dependence of both within-cycle and time-averaged swimming speed on gait dynamics, with the trot out-competing the remaining gaits [10]. A significant difference between the two variants on the gallop (either CW or CCW) was also observed when the appendages are attached in a parallel configuration relative to the body, this introduces a chirality similar to that present in our model.

The performance differences achieved between the gaits can be understood by examining the relative displacement achieved *within* each beat cycle (Fig. 4c). Here, displacement is measured in units of body-lengths/cycle relative to the start position at the beginning of each prospective power stroke of the first (reference) flagellum (*φ*_1_). We observed that the pronk gait achieved the largest positive displacement (forward swimming) during the power stroke, but this progress is negated by a similarly large negative displacement (background) during the recovery stroke. Meanwhile in the trot gait, there is always at least one pair of flagella executing a power stroke at all times, leading to a largely monotonic displacement curve (always forwards), and ultimately the greatest net per-cycle progression. These observations are again in good agreement with the equivalent scenarios measured in the roboflagellate (see for example [10], Fig. 8).

We briefly compare the hydrodynamic efficiencies of the different gaits. We evaluate swimming efficiency using the definition introduced by Lighthill [42, 43], where the power generated by the flagella averaged over a single beat cycle 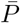 is compared to the external power needed to maintain a rigid sphere in uniform motion with velocity 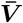 :

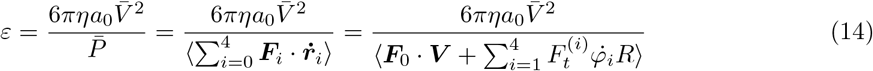

where ⟨ ·⟩denotes a time-average. Compared to the biflagellate case (CR), where *ε*_CR_ = 0.06, we find *ε*_gallop (ccw)_ = 0.07, *ε*_bound_ = 0.09, *ε*_gallop (cw)_ = 0.10, *ε*_pronk_ = 0.09 and *ε*_trot_ = 0.13.

## V. CONFIGURATIONAL ASYMMETRIES

In the previous section, we observed that inherently asymmetric gaits such as the pronk or bound can accentuate asymmetries in the free-swimming trajectories. Further asymmetries can also arise from different sources, including in the respective beat dynamics of the four flagella, or in the geometry of the scaffold.

One possibility is to introduce an asymmetry in the bead rotation (i.e. flagellar beat) frequencies, for example between opposite pairs of flagella. In this scenario however, the phase differences between flagella in a given gait is not preserved. Just as in the biflagellate case, it is likely that real cells tune the balance of flagellar beating to control the overall trajectory heading, particularly in the context of symmetry-breaking and tactic behaviours [27]. A detailed investigation is beyond the scope of the present study.

An alternative is to introduce a geometric asymmetry so that the four flagella beat planes no longer obey the same tilt angle. In the example configuration shown in Fig. 5a,b, the tilt angle *β*_(*ij*)_, each pair of opposite flagellar beads *i, j* move on orbits tilted by an angle *β*_(*ij*)_, with *β*_(13)_ ≠ *β*_(24)_. This quasi-rotational symmetry has been documented in the flagellar apparatuses of some species of green algae [44]. This configuration further breaks axial symmetry and again induces superhelical trajectories in all gaits, including in the previously symmetrical gaits such as the pronk and trot (Figure 5c).

**FIG. 5.**
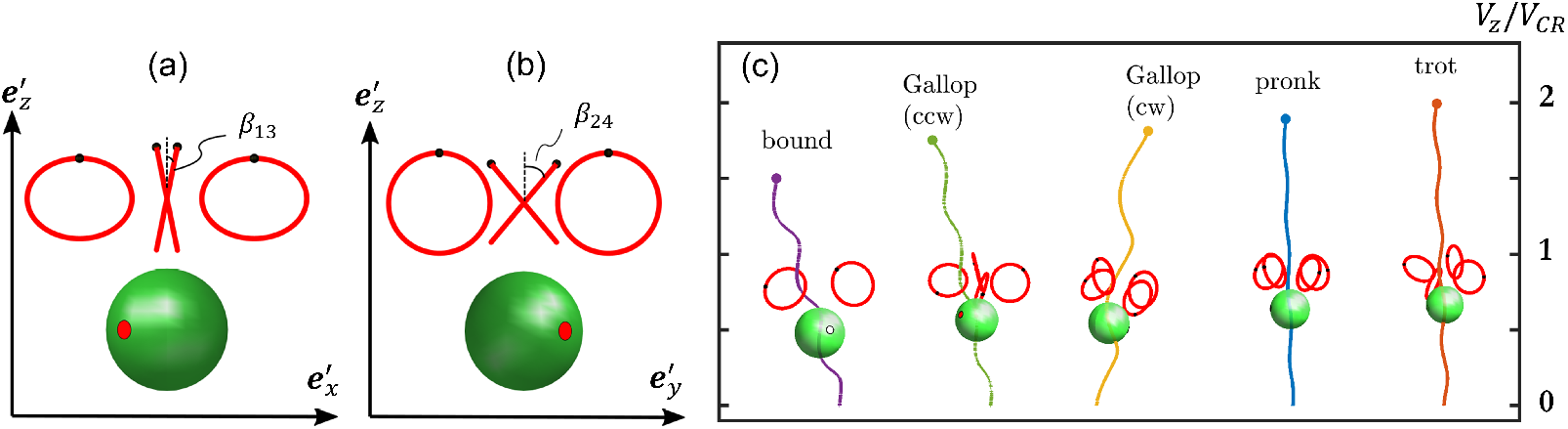
Further geometric asymmetries can also influence the overall symmetry of the trajectory. For instance we can vary the tilt angle so that opposite flagella pairs have the same tilt *β*_(_*ij*), (a) *β*_1_ = *β*_3_ = *β*_(13)_, (b) *β*_2_ = *β*_4_ = *β*_(24_. (c) Sample trajectories for the different gaits, with *β*_(13)_ = 0.3 and *β*_(24)_ = 0.25.

## VI. CLEARANCE RATE, FEEDING FLOWS, AND PREDATION

When considering the ecological or evolutionary drivers of gait diversification in flagellates, it is important to recognise that flagellar actuation strategies that are optimized for swimming may not necessarily be optimal for other important physiological processes, such as feeding [16, 45]. Unlike photoautotrophic algae, some marine flagellates are obligate heterotrophs [46], meaning that they must supplement their diet by phagocytosing bacteria or other small prey organisms - a process that may be significantly influenced by the precise coordination patterns of the flagella. In this section we explore the functional relationships between gait and induced near-field flow architecture in the context of feeding, prey handling and possible evasion from predators.

We quantify the clearance rate of a feeding microorganism as the equivalent volume of water from which the micro-swimmer removes all feed or prey particles per unit time. In suspension-feeding microorganisms, this can be modelled by the filtration rate at which water is passed through a well-defined filtering surface [47]. Here, we model the catchment area of the feeding filtration with a two-dimensional disk Σ of radius *ϱ* located at a distance *H* in front of the cell body. The clearance rate is the volumetric flow rate *Q*_Σ_ across the disk Σ:

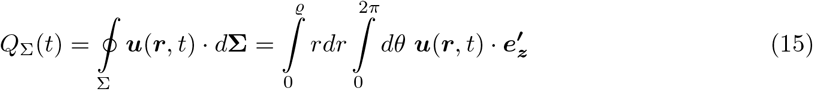

The discretised version of (15) is 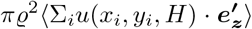, where the index *i* indicates the *i*-th time step and the sum is over an entire beat cycle. Here we chose *ϱ* = *a*_0_, the cell radius.

The average clearance rate over the entire beat cycle 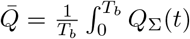 *dt* can be used to quantify the overall effect of the cell on the region described by Σ (Fig. 6a). 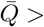 0 for all values of *H*, and for all gaits, showing that flagellar motion has the net effect of pushing fluid away from the front part of the cell body. However, the clearance rate has a maximum at *H ∼* 5.5*µm*, corresponding to where Σ intersects the centers of the flagellar orbits.

**FIG. 6.**
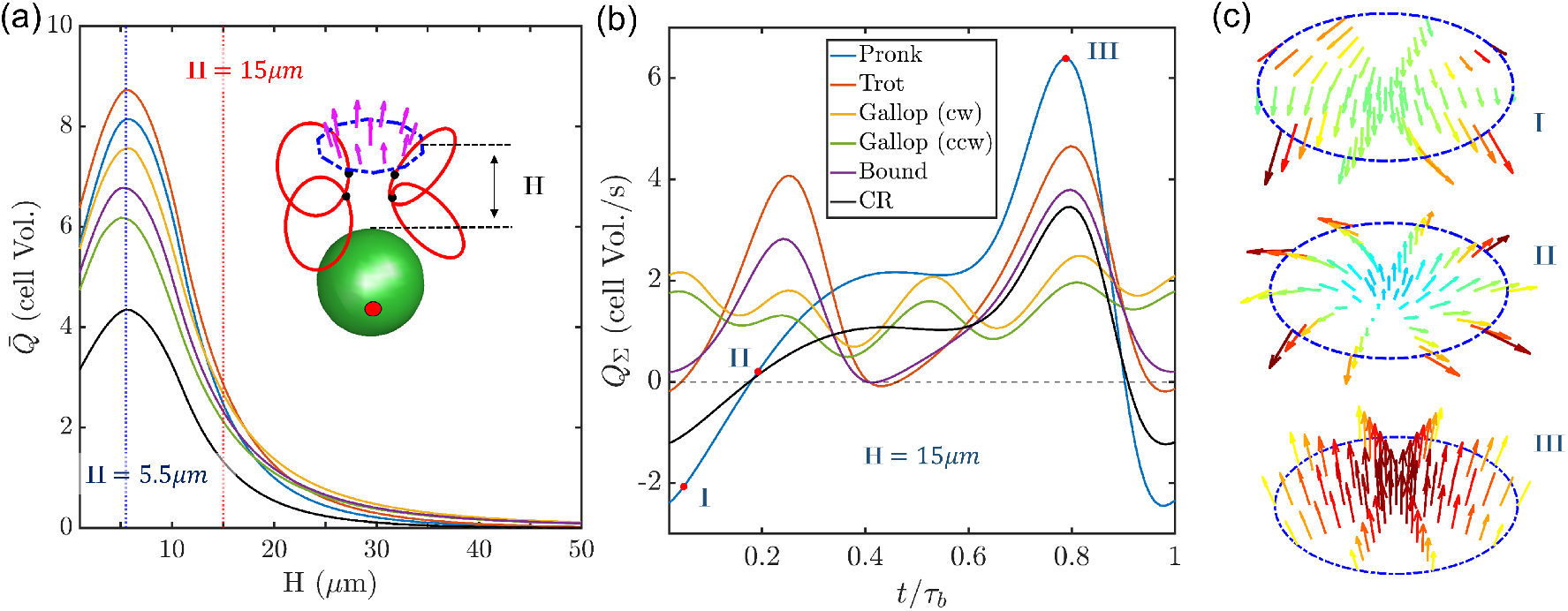
(a) Period-averaged clearance rate as a function of distance from the cell body. Inset: configuration showing the five beads and relative placement of a clearance disk. (b) Within-beat cycle changes in clearance rate again depends on gait. (c) For *H* = 15, snapshots highlight the reversal of flow direction over one cycle for the pronk gait.

Upon analysing how the time-dependent clearance rate *Q*_Σ_ varies throughout the beat cycle (see Fig. 6a), we observe that if the disk is located between the cell body and the tip of the flagellar orbit, *Q*_Σ_ assumes negative values (corresponding to fluid being pulled towards the cell body) only during the recovery stroke for the biflagellate and pronk gaits. Conversely, if the disk is placed in front of the stagnation point, all the gaits except for the two gallops display *Q*_Σ_ *<* 0 during the power stroke, and the beat cycle average is smaller, signifying that a smaller volume of fluid is pushed away from the cell body. In this case, differences between gaits are also more evident, with the pronk gait showing the highest negative clearance rate (see Fig. 6b).

The in-beat differences in *Q*_Σ_ suggest that some gaits can be used to effectively pull particles suspended in the fluid towards the stagnation point, then letting them approach the front of the cell body either by triggering a “stop” response or by exploiting the coasting effect achieved during the *Q*_Σ_ ≤ 0 phase of the beat. The spatial arrangement of flagella forces around a cell is known to greatly influence the statistics of particle entrainment and near-wall contact probabilities [48].

Many flagellates are themselves the prey of other species. The flow disturbances created by any swimmer necessarily dictate the likelihood that they will be perceived by an approaching predator. Finally, we ask whether gait can also influence the hydrodynamic perception field of quadriflagellates, and thereby facilitate quiet-swimming [49].

Fig. 7 shows the spatial decay of the components of the average fluid velocity 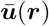, obtained by calculating (12) from the solutions ***r***_*i*_(*t*), and averaging it over a complete beat cycle, with a time step Δ*t* = 0.01. In the far-field, the flows broadly obey the usual stresslet decay (*∼ r*^*−*2^) corresponding to the simple flow field generated by two oppositely directly stokeslets. However it is clear that flow field attenuation is gait-dependent, suggesting that organisms can also manipulate streamlines by changing their swimming gait to avoid predators. For front-mounted swimmers, the difference is most pronounced in the z-direction, aligned with the cell’s anterior-posterior axis, which is likely the direction most sensitive to approaching predators. A rotary gallop gait with a chirality counter to the sense of the individual tilt angle of the flagella, also appears to generate the greatest flow disturbances (e.g. compare the CW with CCW gallop).

**FIG. 7.**
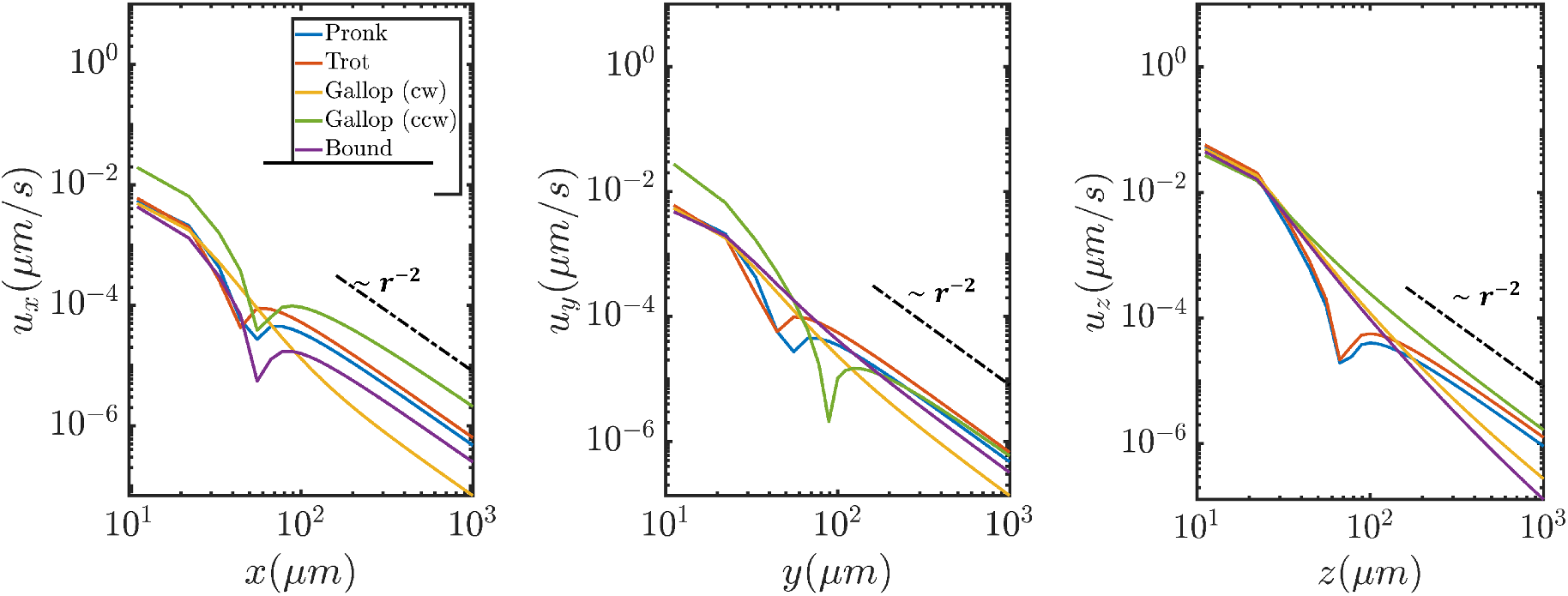
Components of the average fluid flow velocity ***u***(***r***) evaluated for each of the different gaits.

## VII. CONCLUDING REMARKS

In summary, we have presented a novel hydrodynamic model of a prototypical quadriflagellate microswimmer, in which flagellar dynamics are modelled as small beads rotating along circular orbits. By prescribing the action, particularly, the order of actuation of these four flagella, we systematically compared the consequences of gait on the induced flow architecture, free-swimming dynamics and efficiency, as well as for the manipulation of feeding flows. These quantitative comparisons were made against physical measures of flow features including swimming speed, angular rotation, swimming efficiency, clearance rate etc, with particular emphasis on the three-dimensional nature of these flow disturbances. Our approach allowed us to visualise the detailed near and far-field flow patterns around individual swimmers, and also to resolve how these highly-dynamic patterns evolve over time during the course of a stereotypical beat cycle.

So far, we have focused solely on gaits that are stable in time, in which the swimmer dynamics are prescribed. In reality, long-time tracking of single flagellates has shown that gait and behaviour are both highly-dynamic, exhibiting stochastic and reversible fluctuations between distinct gaits or swimming modes [20]. Future work should seek to account for the statistics of such gait transitions as measured from experimental data to investigate their influence on the overall morphology of simulated trajectories.

While it is not yet feasible/possible to directly manipulate the gaits of live cells, our work suggests novel strategies for gait-based control of swimming efficiency in systems of artificial microswimmers. Understanding the detailed three-dimensional distribution of stresses and the time-dependence of induced flows, can help inform designs of artificial devices capable of spatiotemporal flow manipulation at the microscale [50, 51].

Previous experimental studies have already demonstrated the range of possible swimming and foraging activities induced by diverse flagellates with varying numbers, patterns, arrangements and types (e.g. hairy vs naked) of flagella [7, 52]. Our present work reveals that even for organisms exhibiting *identical* flagella number, morphology and beat pattern, distinct propulsion strategies and speeds can still emerge - in other words, gait alone can strongly dictate a cell’s swimming dynamics. This is consistent both with biological measurements from different quadriflagellate species, and also with laboratory measurements conducted with an up-scaled robophysical model [10]. The ‘trot’ gait in particular, was found to be the most efficient of all quadriflagellate gaits. Other gaits all achieved linear and angular speeds that were faster than that of the biflagellate model, to different degrees.

From an eco-evolutionary perspective, the existence of distinct gaits of self-propulsion in morphologically comparable species is consistent with the notion that many flagellates are mixotrophic, meaning that they must acquire resources through consuming prey organisms or particulate matter in the medium, in addition to directed swimming and photosynthesis. These different flagellar actuation gaits may have emerged over the course of evolution in response to adaptation and survival in different habitats, to address a fundamental trade-off between swimming and other key processes not directly motility - e.g. feeding and stealthy living. The distinct spatiotemporal flow signatures produced by each of the gaits also suggests a concrete route by which organisms can manipulate their near-field flows in order to accommodate other specialised motor actions, such as feeding or prey-detection by other sensory appendages [35, 53].

## Supporting information

Imposed bead dynamics corresponding to four quadriflagellate gaits (trot, pronk, gallop (cw), bound). Note the 3D movement of each flagella bead.

Near-field flow fields (side view: x-z plane, and top view: x-y plane) induced by the trot gait.

Near-field flow fields (side view: x-z plane, and top view: x-y plane) induced by the pronk gait.

Near-field flow fields (side view: x-z plane, and top view: x-y plane) induced by the CW gallop gait.

Near-field flow fields (side view: x-z plane, and top view: x-y plane) induced by the bound gait.

## ACKNOWLEDGMENTS

This work received funding from the European Research Council (ERC) under the European Union’s Horizon 2020 research and innovation programme (grant agreement No 853560 *EvoMotion*, to KYW).

## VIII. APPENDIX A

List of supplementary videos

- Video 1: Imposed bead dynamics corresponding to four quadriflagellate gaits (trot, pronk, gallop (cw), bound). Observe the movement of each flagella bead in a tilted plane (*β* ≠ 0).
- Videos 2-5: Near-field, three-dimensional flow fields (side view: x-z plane, and top view: x-y plane) induced by each quadriflagellate gait.

